# *p75NTR* neurotrophin receptor function is redundant for development, growth and fertility in the rat

**DOI:** 10.1101/2023.12.27.573424

**Authors:** Stephen Meek, Karamjit Singh-Dolt, Linda Sutherland, Matthew G.F. Sharp, Jorge Del-Pozo, David Walker, Tom Burdon

## Abstract

The p75NTR neurotrophin receptor has positive and negative roles regulating cell survival in the nervous system. Unambiguous interpretation of p75NTR function *in vivo* has been complicated, however, by residual expression of alternate forms of p75NTR protein in initial *p75NTR* knock-out mouse models. As rats are the preferred rodent for studying brain and behaviour, and to simplify interpretation of the knock-out phenotype, we report here the generation of a mutant rat devoid of the p75NTR protein. TALEN-mediated recombination in embryonic stem cells (ESCs) was used to flank exon 2 of *p75NTR* with Lox P sites and produce transgenic rats carrying either un-recombined floxed *p75NTR^Ex^*^2*-fl*^, or recombined, exon-2 deleted *p75NTR^Ex^*^2*-*^*^Δ^* alleles. Crossing *p75NTR^Ex^*^2*-fl*^ rats with a *Cre*-deleter strain efficiently removed exon 2 *in vivo*. Excision of exon 2 causes a frameshift after p75NTR Gly23 and eliminated p75NTR protein expression. Rats lacking p75NTR were healthy, fertile, and histological analysis did not reveal significant changes in cellular density or overall structure in their brains. Thus, p75NTR function appears largely dispensable for normal development, growth and basal homeostasis in the rat. The availability of constitutive and conditional *p75NTR^Ex2-Δ^* rats should, however, provide new opportunities to investigate specific roles of p75NTR upon injury and during regeneration.

## Introduction

The p75NTR receptor is a key mediator of neurotrophin signalling and has been ascribed a variety of roles in regulating the growth and maintenance of the nervous system ^1,2^. The founding member of the tumour necrosis factor receptor family, p75NTR is a single membrane spanning protein that lacks intrinsic catalytic activity but contains an intracellular death domain ^3^. Depending on its interacting partners p75NTR can either promote neuron outgrowth and survival, or induce cell death ^2^. In association with the neurotrophin Trk receptors, p75NTR forms a high affinity receptor for mature processed neurotrophins and promotes cell survival. By contrast, when linked to the sortilin receptor and bound by unprocessed proneurotrophin ligands, p75NTR induces apoptosis. When bound to the Nogo receptor, p75NTR mediates signalling downstream of myelin proteins which control cell growth and resolution of immune responses. Through these diverse interactions p75NTR is thought to influence the balance between cell growth, survival and death within the nervous system ^2^.

Elucidating the functional role of p75NTR *in vivo* has relied upon studies of a series of knock- out mice. Mice homozygous for the deletion of exon 3 are viable and fertile, but have impaired innervation of peripheral sensory neurons, and reduced response to the neurotrophin NGF ^4^. Subsequent studies, however, showed that mice harbouring the exon 3 deletion still express a short form of p75NTR, complicating conclusions regarding p75NTR function ^5^. Disruption of *p75NTR* exon 4 eliminated expression of both the long and short from receptors in mice, but also generated a cryptic protein containing just the death domain ^6^. This likely contributed to the high mortality of pre and post-natal pups, and growth retardation of surviving *p75NTR* exon 4 disrupted mice. Indeed, more recently, *p75NTR* knock-out mice carrying a deletion of exons 4-6 encoding the transmembrane and cytoplasmic domains, lacked any p75NTR functional protein but were viable, reproduced normally and displayed only relatively subtle neurological defects ^7^. This suggests that, in contrast to early reports and despite the involvement of p75NTR in several signalling pathways and cellular processes, p75NTR function might be largely redundant in the mouse.

To evaluate p75NTR function in another mammal, and as a comparison with the phenotypes described in previous mouse studies, we generated knock-out rats lacking p75NTR receptor. The laboratory rat is a preferred animal model in studies of brain function and the nervous system, due to its larger brain size, greater intelligence and mental flexibility when compared with the mouse ^8,9^. Until relatively recently gene targeting in the rat was not straightforward. However, the development of genuine germ line competent rat ESCs and the advent of *in ovo* gene editing using programmable nucleases have enabled the routine application of state-of-the-art genetic engineering techniques to the rat ^10^. In this report we describe using ESCs to generate mutant rats that are constitutively or conditionally deficient for p75NTR, providing a new animal model with which to dissect the apparently contrasting functions of this enigmatic neurotrophin receptor.

## Results

### Generation of *p75NTR^Ex^*^2^*^-Δ^* and *p75NTR^Ex^*^2^*^-fl^* alleles in rat

We used targeted homologous recombination in ESCs to generate constitutive and conditional mutations in the rat *p75NTR* gene. Guided by a design strategy recommended by the EUCOMM mouse gene targeting consortium (https://www.mousephenotype.org/about-impc/about-ikmc/eucomm/) we elected to delete exon 2 of rat *p75NTR*, which results in a frameshift and the production of a 31 amino acid polypeptide containing only the first 23 amino acids of p75NTR (Figure 1A). We generated a TALEN gene editing nuclease designed to introduce a double strand DNA break within intron 1, 393 bp upstream of exon 2; to promote the insertion of a neomycin resistance cassette nested by two Frt recombination sites, and flank exon 2 with two LoxP recombination sites (Figure 1B). DAK31 rat ESCs ^11^ were electroporated with plasmids encoding the TALEN and one of two neomycin resistant targeting vectors, containing either a floxed exon 2 or the *in vitro* Cre-mediated recombinant deleted for exon 2. After 48 hours recovery, the electroporated cultures were subjected to selection in 200 μg/ml G418 for a further 10 days. The resulting neomycin resistant rESC colonies were picked, expanded and screened by PCR and Southern Blotting for correct insertion of the targeting construct (Figure 1C). Correctly targeted clones were transiently transfected with plasmid expressing FLP recombinase and subclones in which the neomycin cassette had been excised were selected based on sensitivity to neomycin. *p75NTR* exon 2 deletion subclones B10 and D10, and the floxed *p75NTR* subclone 34.4, having a normal karyotype and developmental potential as judged by embryoid body differentiation, were injected into Sprague Dawley (SD) blastocysts to generate ESC-derived chimaeric rats. Male chimaeric rats were crossed with SD females and germ line transmission of mutant *p75NTR* alleles through their offspring was used to establish transgenic lines of rats carrying either the *p75NTR* exon 2 deletion (*p75NTR^Ex^*^2^*^-Δ^*), or the floxed *p75NTR* (*p75NTR^Ex^*^2^*^-fl^*) allele (Table 1). Crossing *p75NTR^Ex2-Δ^* heterozygous rats generated homozygotes that were viable and born at a normal Mendelian frequency (Table 2). To confirm that the *p75NTR* ^Ex2-fl^ allele could be efficiently recombined *in vivo*, male *p75NTR^Ex2-fl/fl^* rats were bred with female *Cre*-deleter *CAGGS-Cre* transgenic rats ^12^. PCR analysis of resulting embryos showed that exon 2 was efficiently deleted in all embryos by the Cre activity present within all the developing CAG-Cre oocytes (Figure 2A-C) ^13^.

**Figure 1.**
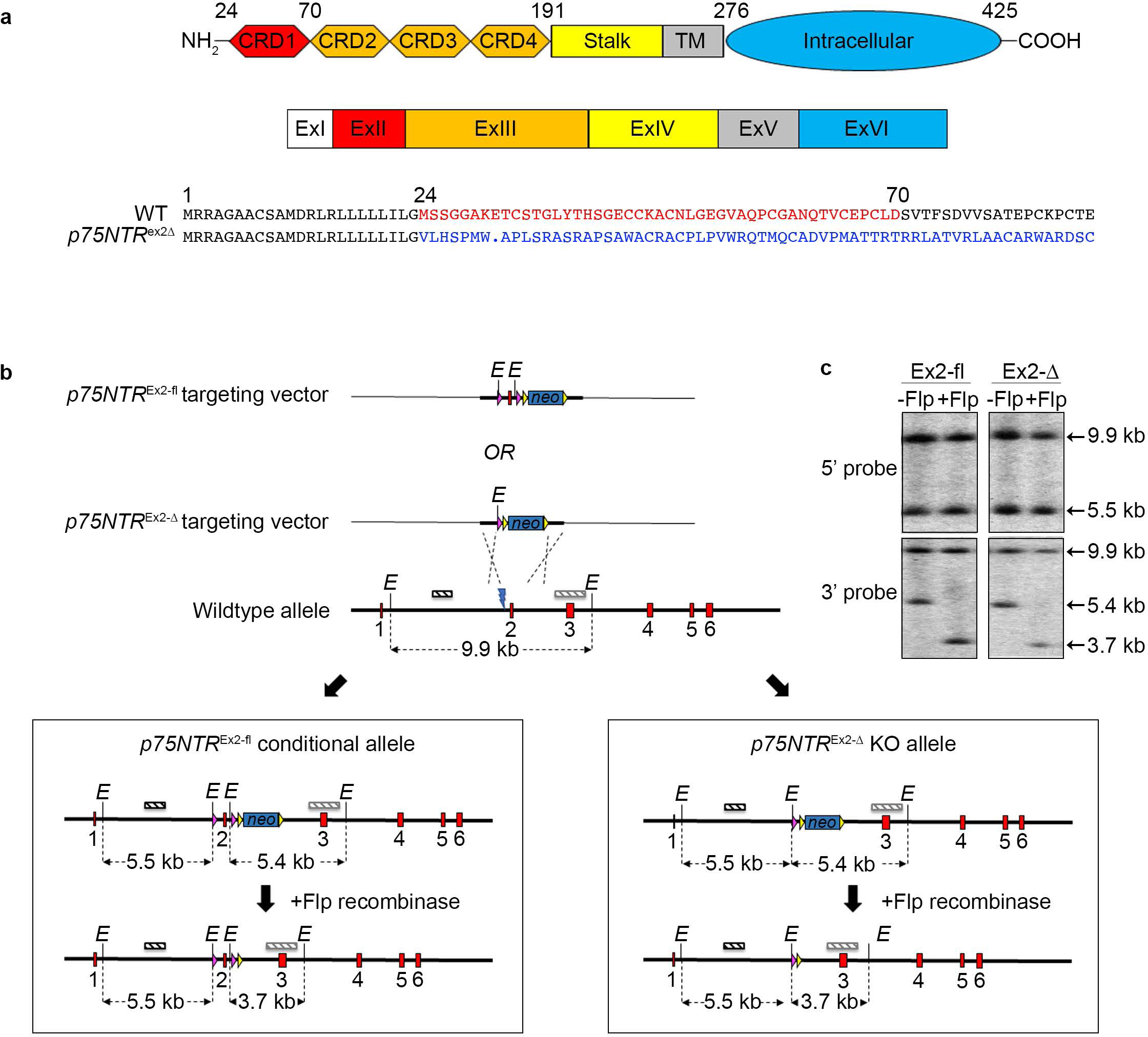
Generation of KO and conditional KO *P75NTR* rats. (a) The domain structure of p75NTR protein depicting the N-terminal extracellular domains consisting of four cysteine-rich domains (CRD1-4) and stalk region, the transmembrane domain (TM) and the intracellular C- terminal domain that contains the Chopper and Death domains (top schematic). The domains are encoded by six corresponding coding exons (middle schematic). Deletion of exon 2 (red) is predicted to generate an out-of-frame mutation at 24aa (blue sequence) and introduce a premature STOP codon at 31aa (blue dot) (bottom schematic). (b) Schematic of the *p75NTR* KO strategy showing the *p75NTR^Ex2-fl^*and *p75NTR^Ex2-Δ^* targeting vectors, the wild-type *p75NTR* allele and the *p75NTR^Ex2-fl^* and *p75NTR^Ex2-Δ^* alleles before and after FLP-mediated recombinase removal of the selection cassette. LoxP sites (pink arrowheads) flank exon 2, 408 bp 5’ and 325 bp 3’, with the FRT-flanked PGKneo selection cassette (blue box flanked by yellow arrowheads) 3’ of the 3’ loxp site. TALEN-mediated double-strand break (blue lightning bolt) was designed to promote homology-directed repair. Exons are depicted by red boxes, non-exonic genomic DNA and plasmid DNA by thick and thin black lines respectively. The EcoRI (E) restriction site and 5’ and 3’ probe sequences (dark and light hashed boxes respectively) used for Southern blot screening, and expected restriction digest fragments sizes are shown. (c) Southern blot analysis of EcoRI-digested genomic DNA from *p75NTR^Ex2-fl^* and *p75NTR^Ex2-Δ^* targeted ESCs with (-Flp) and without (+Flp) PGKneo selection cassette, using 5’ and 3’ external probes (top and bottom panels respectively).

**Figure 2.**
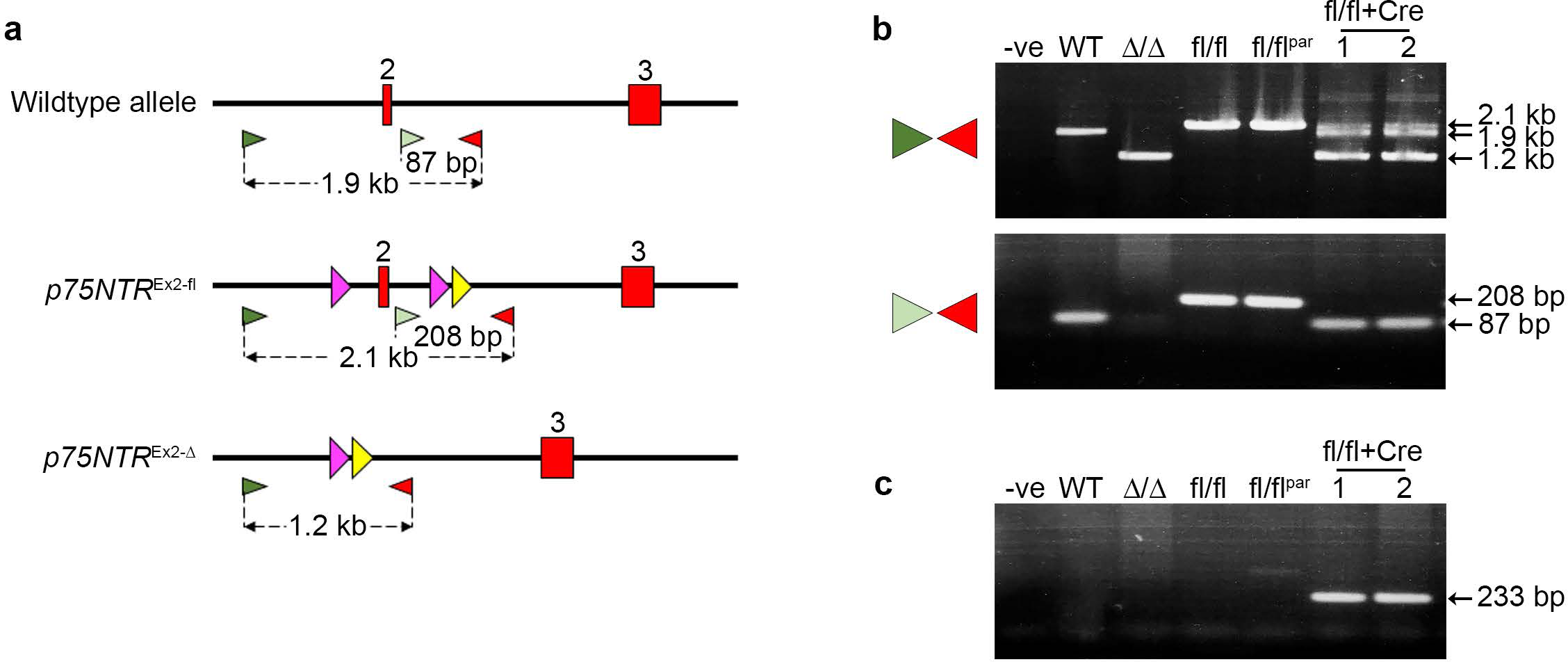
In vivo Cre recombinase-mediated deletion of conditional *p75NTR* allele. (a) PCR screening strategy used for detecting Cre recombinase-mediated deletion of exon 2 and discriminating between wild-type, *p75NTR^Ex2-fl^* and *p75NTR^Ex2-Δ^* alleles. The locations of exons (red boxes), non-exonic genomic DNA (thick black lines), loxP sites (pink arrowheads), FRT sites (yellow arrowheads), forward PCR primers (light and dark green arrowheads) and reverse PCR primers (red arrowheads) are shown. Sizes of expected PCR fragments are highlighted. (b) PCR analysis of genomic DNA from wild-type rat ESCs (WT), and *p75NTR^Ex2-Δ/Δ^ (Δ/Δ)* and *p75NTR^Ex2-fl/fl^ (fl/fl)* homozygous rats along with offspring (1 & 2) generated from breeding of a male homozygous *p75NTR^Ex2-fl/fl^*(fl/flpar) with a female heterozygous Cre transgenic rat. The upper panel shows amplicons generated using the 5’ forward primer (dark green arrowhead) and reverse primer, demonstrating deletion of Exon 2 in *p75NTR^Ex2-fl/^* /CRE offspring 1 &2). The lower panel shows amplicons generated using the 3’ forward primer within the deleted region 5’ to exon 2 (lower panel) and the reverse primer. (c) PCR amplification of a 233 bp DNA fragment from the samples in 2b using primers specific for the CRE transgene.

**Table 1.** Germline transmission efficiency.

| Cell line | Genomic modification | Chimaeras (sex) | GLT |
| --- | --- | --- | --- |
| 34.4 | $p75NTR^{ex2fl}$ | 7 (6M/1F) | 4 (57%) |
| B10 | $p75NTR^{ex2\Delta}$ | 14 (14M) | 1 (7%) |
| D10 | $p75NTR^{ex2\Delta}$ | 6 (6M) | 2 (33%) |

**Table 2.** Viability of P75NTR KO rats.

| Genotype | Number of offspring* (% of total) | Sex |
| --- | --- | --- |
| Wildtype | 9 (18%) | 4M + 5F |
| Heterozygous | 30 (59%) | 11M + 19F |
| Homozygous | 12 (23%) | 6M + 6F |

\* Offspring from seven litters of a GLT x GLT mating.

### Deletion of *e*xon 2 eliminates p75NTR protein expression

To determine how deletion of exon 2 affected p75NTR expression we crossed heterozygous *p75NTR^Ex2-Δ/+^* rats and analysed brain tissue of their offspring. *p75NTR* RNA expression in *p75NTR^Ex2-Δ/+^* heterozygotes was readily detected by RT-qPCR using primers annealing downstream of the exon 2, in exons 5 and 6. By contrast, the exon 2-deleted RNA was almost completely absent in homozygous *p75NTR^Ex2-Δ/Δ^* rats suggesting that premature translation termination might destabilise the mutant RNA (Figure 3A). Analysis of p75NTR protein expression in corresponding tissue lysates by Western blotting, using a polyclonal antibody recognising the intracellular domain of p75NTR, did not detect any p75NTR proteins in the brain of *p75NTR^Ex2-Δ/Δ^* rats (Figure 3B). By contrast, *p75NTR^Ex2-fl/fl^*rats expressed p75NTR proteins at levels equivalent to that in wild-type *p75NTR* rats (Figure 3C). We conclude that deletion of *p75NTR* exon 2 eliminates p75NTR protein, generating a functionally null allele, whilst in the floxed *p75NTR^Ex2-fl/fl^* rats, p75NTR expression was unaffected.

**Figure 3.**
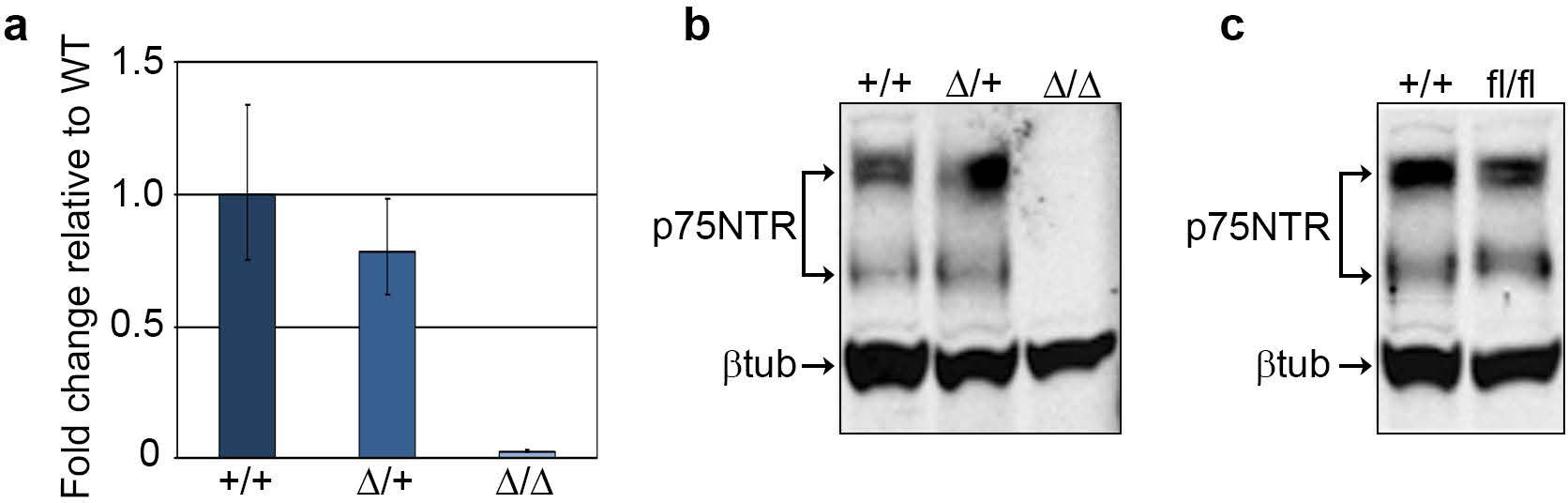
Loss of *p75NTR* expression in KO rats. (a) RT-quantitative PCR analysis of *p75NTR* in wildtype (+/+), heterozygous (Δ/+) and homozygous (Δ/Δ) *p75NTR^Ex2-Δ^* rat brains (mean ± sd of three technical replicates). (b) Western blot analysis of p75NTR protein expression in wildtype (+/+), heterozygous (Δ/+) and homozygous (Δ/Δ) *p75NTR^Ex2-Δ^*rat brains. (c) Western blot analysis of p75NTR protein expression in wildtype (+/+), and homozygous (fl/fl) *p75NTR^Ex2-fl^* rat brains.

### Rats lacking p75NTR are healthy and fertile

p75NTR-deficient rats were visually indistinguishable from their p75NTR expressing littermates: they exhibited similar caged behaviour and possessed normal fertility. Indeed, mating p75NTR-deficient rats produced average sized litters of healthy pups (Table 3). As p75NTR-deficient mice have been reported to have deficits in neuronal development, we compared the histology of brains of *p75NTR^Ex2-Δ/Δ^* and *p75NTR^Ex2-fl/fl^* rats. Brain cellular density and structure in cerebellum, caudoputamen and thalamus regions was indistinguishable between the p75NTR expressing and deficient rats (Figure 4A, C and Supplementary Table S1). It was noted that brain size (weight) when normalised relative to body weight, was increased slightly in p75NTR-deficient rats (Figure 4B). Other organs examined did not show this difference (Supplementary Table S2). Based on this general assessment, we conclude that under normal circumstances p75NTR has non-essential roles during development, growth and basal homeostasis of the adult rat.

**Figure 4.**
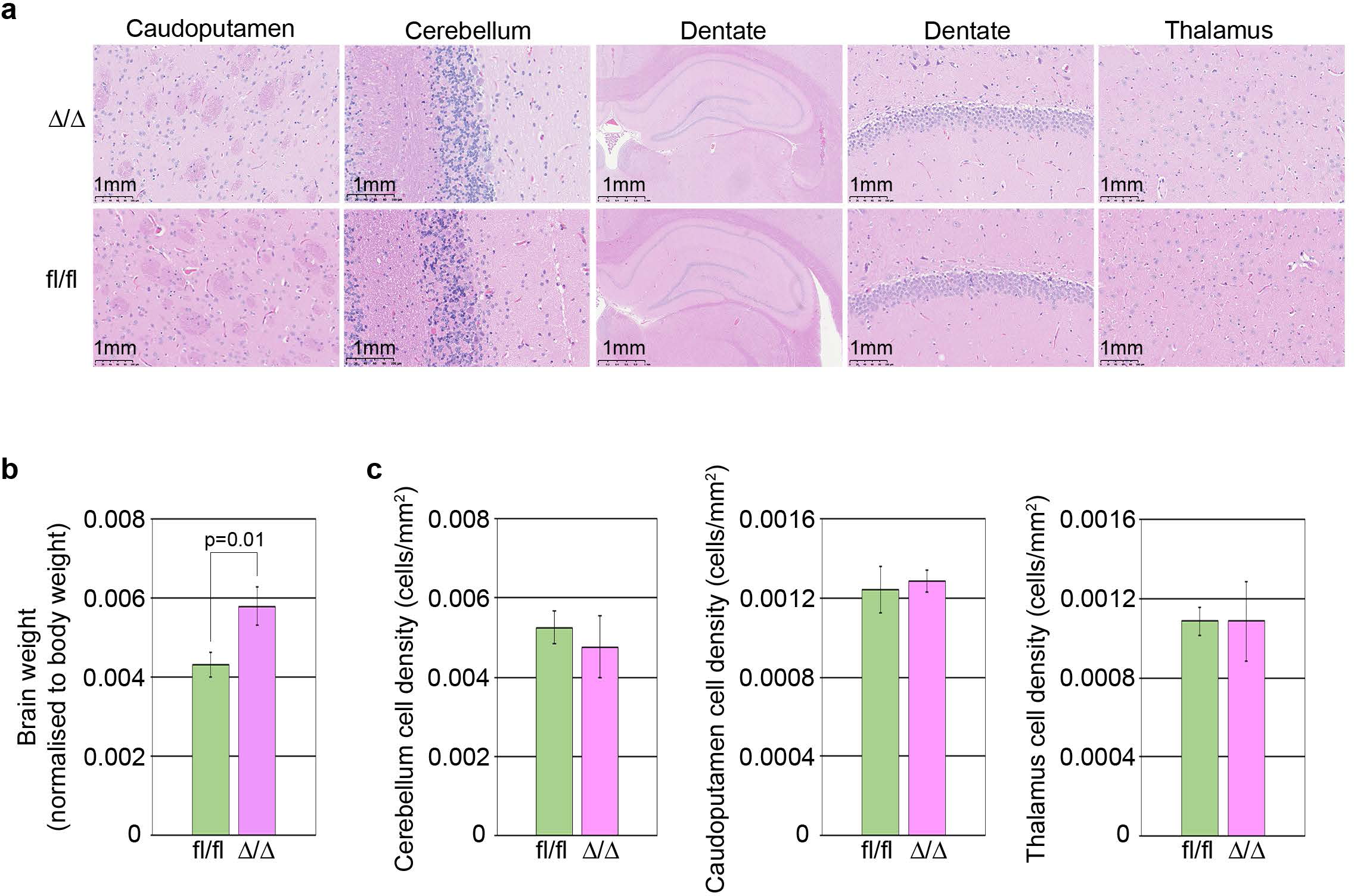
Histopathological analysis of *p75NTR^Ex2-Δ/Δ^* and *p75NTR^Ex2-fl/fl^* rats. (a) Representative images of haematoxylin and eosin stained histological sections from male *p75NTR^Ex2-fl/fl^* and *p75NTR^Ex2-Δ/Δ^* homozygous rats. (b) Whole, fixed brain weight normalised to body weight, for *p75NTR^Ex2-fl/fl^* and *p75NTR^Ex2-Δ/Δ^* homozygous rats. The statistical analysis was performed using an unpaired two sample t-test with a 5% significance level (mean ± sd of three biological replicates). (c) Cell density in the cerebellum (left panel), caudoputamen (middle panel) and thalamus (right panel) of *p75NTR^Ex2-fl/fl^* and *p75NTR^Ex2-Δ/Δ^* homozygous rats (mean ± sd of three biological replicates).

**Table 3.** Fertility of P75NTR KO rats.

| KO x KO | Litter size |
| --- | --- |
| Breeding 1 | 10 (5M + 5F) |
| Breeding 2 | 7 (4M + 3F) |
| **Average litter size** | **8.5 (4.5M + 4F)** |

## Discussion

The integration of neurotrophin-mediated cell survival and death signals by p75NTR is regarded as an important regulator of the fate of neurons and other cell types. However, the complexity of hypomorphic or gain–of-function mutations associated with the initial p75NTR transgenic studies in mice, upon which many early conclusions were based, can defy simple interpretation ^4,5^. By contrast, constitutive and conditional deletion of *p75NTR* exon 2 generates a complete null allele in the rat and secures a simple unambiguous experimental platform for exploring the function of p75NTR *in vivo*. Indeed, the viability and apparent health of the p75NTR-deficient rats prompts further examination of the role of p75NTR in neurotrophin signalling out-with normal development and basal homeostasis.

Deletion of *p75NTR* exon 2 eliminates expression of p75NTR protein in the rat. The mutation causes a translation frameshift that generates the first 23 amino acids of the N-terminal signal peptide of p75NTR ^14,15^, and an additional 7 unrelated amino acids. The mutation also suppresses expression of the *p75NTR^Ex2-Δ^* transcript, presumably through a nonsense-mediated decay mechanism ^16^ . Two different *p75NTR* mutant lines of mice are reported to completely eliminate p75NTR protein expression. Bogenmann and colleagues produced transgenic mice in which the region encompassing exons 4-6 is flanked by LoxP sites (*p75NTR^FX^*), thus allowing Cre-mediated deletion of the majority of the p75NTR intracellular domain, either constitutively or conditionally when crossed with suitable *Cre-*deleter strains of mice ^7^. By contrast, Boskovich and colleagues, inserted a LoxP recombination cassette in the exon1/intron 1 region (*p75NTR^Inv^*), that inverts its orientation through the action of Cre, replacing the coding region of Exon 1 with a mCherry reporter ^17^. Both the *p75NTR^FX^*and *p75NTR^Inv^* alleles enable the selective deletion of p75NTR from specific neuronal cells in mice. In a similar way selective deletion of rat p75NTR can be now achieved in rats carrying the *p75NTR^Ex^*^2^*^-fl^* allele, either by crossing with Cre-driver strains, or by delivering Cre using viral vectors.

Given some of the phenotypes attributed to the loss of p75NTR in mice, we were surprised to find that p75NTR-deficient rats developed normally, were fertile and displayed no overt differences in growth or general behaviour from control rats. Although the mixed genetic background of the p75NTR-deficient rats (DA and Sprague Dawley) might mask subtle phenotypic changes in mutant animals when compared with p75NTR-expressing littermates, the roles of p75NTR in normal development and basal homeostasis appear to be non-essential in the rat. Interestingly, although p75NTR-deficient mice carrying a deletion of exons 4-6 were reported to exhibit minor signs of peripheral neuropathy, they otherwise appeared relatively normal and were fertile ^7^. The larger size of the brains of *p75NTR^Ex2-Δ^* rats could hint that the normal pattern of neural development is disturbed by loss of p75NTR in rat. However, further experiments are required to eliminate differences that might arise from the slight age difference between the transgenic p75NTR rats in this study. Increases in the numbers of cholinergic neurons have been reported for both *p75NTR* exon 3 and exon 4 deleted mice ^18^. Moreover, an increased size of the cerebellum was described in p75NTR mutant mice and was thought to result from over proliferation of cerebellar granule precursor cells ^19^. p75NTR- deficient cerebellar cell cultures derived from a commercially available *p75NTR* knock-out rat (SAGE) also displayed a shortened cell cycle, but no associated effect on cerebellum or brain development was reported ^20^. By contrast, conditional deletion of p75NTR in neural precursor cells of mice, using *Nestin*-driven Cre-mediated inactivation of the floxed *p75NTR^int^* allele, was associated with a reduction in relative brain size and overall size of the mutant mice ^21^. Tissue-specific deletion of *p75NTR* in some cases has been reported to cause a more marked phenotype than constitutive deletion, implying that mechanisms induced early in development may compensate for loss of p75NTR ^19,21^. Future experiments using targeted conditional deletion of the *p75NTR^Ex2-fl^* allele, in conjunction with the corresponding constitutive null allele, should provide an opportunity to explore more precisely the role of p75NTR in the brain and other organ systems of the rat.

The larger size and intelligence of the rat, make this rodent the preferred experimental animal in many studies, particularly when investigating the brain and behaviour ^9^. In addition, the rat has been used extensively as the standard model in pharmacology and toxicology studies. In this respect, the viability of the *p75NTR^Ex2-Δ^* rat provides a simple p75NTR-deficient animal model in which to definitively assess drug action or other experimental interventions. Since many reports implicate p75NTR in regulating responses to injury and regeneration ^22^, the availability of the *p75NTR^Ex2-Δ^* mutations make it feasible now to complement studies in mice, and assess how therapeutic strategies can be applied to ameliorate damage and promote repair, and determine their impact on behaviour and higher cognitive functions.

## Methods

### ESC culture

Rat ESCs (line DAK31), derived from E4.5 rat blastocysts of the inbred pigmented rat strain Dark Agouti were kindly supplied by Professor Austin Smith ^11^. ESCs were maintained on irradiated OF1 or DR4 mouse fibroblasts in 2iL (N2B27 medium, 1 μM PD0325901 [PD], 3 μM CHIR99021 [CH], 1000 U/ml mouse LIF). Cells were passaged every 2–3 days using TVP (0.025 % trypsin, 1 % chicken serum, and 1 μM EDTA) and plated at a density of 0.5–1 x 10^5^/cm^2^. MEK (PD) and GSK3 (CH) inhibitors were supplied by Axon Medchem.

### BAC recombineering

The targeting vector was designed to introduce two loxP sites flanking exon 2 to generate a Cre recombination-mediated conditional knockout of the *p75NTR* gene. The targeting vector was constructed by BAC recombineering using the pSIM17 plasmid as previously described elsewhere ^23,24^. Briefly, the pSIM17 plasmid was electroporated into E. coli cells containing the rat *p75NTR* BAC to facilitate subsequent recombineering steps. First, a single loxP site containing an EcoRI restriction site was introduced in intron1, 393 bp upstream of exon 2. Next, a loxP-frt-PGK-EM7-Neo-frt cassette was introduced into intron 2, 301 bp downstream of exon 2. Finally, the targeting construct consisting of 1 kb homology arms with the loxP flanked exon 2 and frt-flanked neo selection cassette was retrieved into plasmid PL611. The targeting vector was designed so that the TALEN heterodimer binding sites would be disrupted following successful targeting thus preventing cutting of the targeted allele.

### TALEN Construction

The P75NTR TALEN pair (left target – GCAGGTGTGAGGTTTG, right target - CTGGCCCCCTTGGCTAGG) were designed to cut within the spacer region (GGCCAGATTTCAGCTCCA) 393 bp upstream of exon 2, and result in a double strand break adjacent to the loxP insertion site in intron 1. All TALENs were designed using the TALE-NT software (https://tale-nt.cac.cornell.edu/node/add/talen) and assembled by Golden Gate cloning using methods described by Sakuma and co-workers ^25^ . The TALENs were cloned into pCAG-T7 TALEN (Sangamo)-FokI-ELD-Destination and pCAG-T7-TALEN (Sangamo)- FokI-KKR-Destination expression plasmids.

### Gene targeting by homologous recombination

1x10^6^ rat ESCs were plated on to a 6-well plate of mitotically-inactivated mouse DR4 fibroblasts and transfected with 500 ng linearized targeting construct and 250 ng each of left and right TALEN expression plasmids using Lipofectamine LTX (6.75 μl LTX + 2.25 μl PLUS reagent, Invitrogen) according to manufacturer’s instructions. 48 h post-transfection cells were re-plated on to DR4 fibroblasts at a variety of densities ranging from 0.5-10x10^2^ / cm^2^ in 2iL with 150 μg/ml G418 and fed every two days. Resistant colonies were picked between 9-12 days post-plating and expanded for PCR screening and cryopreservation.

### FLP-mediated excision of the neomycin selection cassette

Rat ESCs were transfected as described above with 1 μg pCAGGS-FLP-IP plasmid using Lipofectamine LTX. 48 h post-transfection cells were re-plated on to DR4 fibroblasts at 0.5- 10x10^2^ / cm^2^ in 2iL and fed every two days. Twelve colonies were picked 9-12 d post-plating and expanded.

### Genomic DNA

Cells and tissue were lysed in 500 μl lysis buffer (100mM Tris pH8.0, 200mM NaCl, 5mM EDTA, 0.2% SDS) containing 50 μg proteinase K at 55 °C for at least 2 h. Genomic DNA was precipitated by adding 500 μl isopropanol to the lysate and inverting several times to mix. DNA was pelleted at 9000 g for 10 minutes and the pellet washed twice with 70 % ethanol before air-drying and resuspending in nuclease-free water.

### Southern blot

Eight to ten micrograms of genomic DNA were digested with 40 units of *Eco*RI restriction enzyme overnight at 37 °C. The following morning a further 40 units of enzyme was added and incubated at 37 °C for an additional 5 hours. The resulting DNA fragments were resolved on a 0.7% TAE agarose gel overnight at 25 V. The DNA fragments were UV-nicked prior to transfer to Hybond N + Nylon membrane (GE Healthcare, RPN203B) as described in the manufacturer’s instructions. Following transfer, the DNA was UV cross-linked on to the membrane. Probes were prepared by PCR amplification of the rat *p75NTR* sequence flanking the homology arms to generate a 1004 bp 5’ probe (forward primer TCGTTGCGTTAGTCTTCCTTC, reverse primer GCCTGCTCACTTCTTTTAGGA) and a 1498 bp 3’ probe (forward primer GGCCATTCTGTCCTTTGCCTC, reverse primer GAACCCACCTAGCTCTCAGTA). 25 ng of probe DNA was radioactively labelled with α– dCTP ^32^P using High Prime (Roche, 11 585 592 001), then hybridised to the membrane overnight at 65 °C in Church solution containing 10 μg/ml sonicated Herring Sperm DNA and 10 μg/ml tRNA. Non-specifically bound probe was removed by washing in 2x SSC/0.1 % (w/v) SDS at 65 °C. The membrane was exposed to Kodak Biomax MS film at -80 °C.

### Genotyping

Two independent PCR reactions were performed using 150 ng of genomic DNA. The first reaction consisted of primers designed to flank exon 2 (intr1_F1– AAGGTCTGCCTGTCTAATCTGG, intr2_R1-CATTGATCTTGCAGCACGGAA) to amplify 1959 bp, 2168 bp and 1260 bp from *p75NTR* wild-type, floxed *p75NTR^Ex2-fl^* and the KO *p75NTR^Ex2-Δ^*alleles respectively. The second reaction utilized the same reverse primer and the forward primer (intr2_F1–GAGGTTGGTAGAAGAGCATAGC) located within the Cre-deleted region, 5’ of exon 2 to amplify 87 bp and 208 bp from wild-type and the floxed *p75NTR^Ex2-fl^* alleles respectively. No product is amplified from the *p75NTR^Ex2-Δ^* allele in the second reaction because the 5’ forward primer site is deleted by Cre recombinase-mediated excision of exon 2. Both reactions were performed using NEB Q5 HotStart Taq Polymerase under the following conditions; 98 °C for 1 minute, followed by 32 cycles of 98 °C for 10 s, 67 °C for 30 s and 72 °C for 2 minute with a final extension of 72 °C for 10 minutes. Products were visualised with ethidium bromide on a 1 % (for intr1_F1/ intr2_R1) or 2 % (intr2_F1/ intr2_R1) TAE agarose gel.

### *In vivo* Cre recombination

A homozygous *p75NTR^ex2-fl/fl^* conditional floxed male rat was crossed with a heterozygous female rat carrying the CAG-NCre transgene (Wistar-Tg(CAG-Ncre) 81Jmsk rat strain #301 supplied by RRRC). Skin samples were collected from the head and back regions of 3 day old pups and processed for genomic DNA as described above. Genomic DNA was amplified using primers (forward primer Cre5 - GCGGCATGGTGCAAGTTGAAT and reverse primer Cre3 – CGTTCACCGGCATCAACGTTT) designed to amplify a 233 bp product from the Cre transgene. PCR reactions were performed as above and products visualised with ethidium bromide on a 2 % TAE agarose gel.

### RNA/cDNA preparation

The brains of E13.5 rats were collected in RNALater Stabilisation Solution (ThermoFisher, #AM7020) prior to lysis. RNA was purified using the RNeasy miniprep kit (Qiagen, #74104) according to the manufacturer’s instructions. The recommended on-column DNA digestion was performed and the RNA eluted using 35 μl nuclease-free water. 1 μg RNA was used to synthesise cDNA using the Multi-temp cDNA Synthesis kit (Agilent, #200436). The final reaction volume was made up to 1ml with nuclease free water and used for RT-qPCR analysis.

### RT-qPCR analysis

RT-qPCR analysis was performed using the Platinum SYBR Green QPCR kit (Invitrogen #11744100) containing 10 μl cDNA with forward and reverse primers that hybridised to *p75NTR* exons 5 and 6 respectively (forward primer GAGAAACTGCACAGCGACAG and reverse primer CTCTACCTCCTCACGCTTGG). The reaction was performed on a Stratagene Mx3000 real-time PCR machine under the following conditions; 95 °C for 2 minutes followed by 40 cycles of 95 °C for 15 seconds, then 60 °C for 30 seconds, with a final, melting curve cycle consisting of 95 °C for 1 minute, 60 °C for 30 seconds and 95 °C for 15 seconds.

### Western blot analysis

The brains of E13.5 rats were lysed and 30 μg of protein combined with 5 μl 4x LDS Sample Buffer (ThermoFisher, #NP0004) and 2 μl 10x Sample Reducing Agent (ThermoFisher, #NP0007) in a final volume of 20 μl with water. Samples were heated at 70 °C for 10 minutes prior to loading and electrophoresis on a 10% Bis-Tris polyacrylamide gel (ThermoFisher, #NP0316BOX), followed by electroblotting on to a PVDF membrane. After blocking at room temperature for 1 h with Intercept (PBS) Blocking Buffer (LI-COR, #927-70001) the membrane was probed with anti-p75NTR antibody diluted 1:4000 (Merck, #07-476) and anti-tubulinβ3 antibody diluted 1:500 (Biolegend, #MMS-435P) at 4 °C overnight. The membrane was then probed using the secondary antibodies IRDye680RD (Donkey anti-mouse, 1:5000, Li-Cor, #926-68072) AND IRDye800W (Goat anti-rabbit, 1:5000, LI-COR, #926-32211) at room temperature for 1 h. The membrane was analysed using the LI-COR Odyssey Infrared Imaging System.

### Histological analysis

Brains from the rats were fixed in 4 % neutral buffered formalin. Fixed brains were step-sectioned at the approximate same level for each animal, and embedded in paraffin wax. Sections were stained with haematoxylin and eosin (HE). The histological sections were scanned using the NanoZoomer-XR system (Hamamatsu, Welwyn Garden City, Hertfordshire, UK) to create whole slide digital images. Representative brain areas of interest from *p75NTR^ex2-Δ/Δ^* and *p75NTR^ex2-fl/fl^*males which included cerebellum (white matter, molecular and granular layers), caudoputamen and thalamus, were manually delineated using the QuPath open source software ^26^. Total cell counts of the selected areas were obtained using automated nucleus detection, and the total area calculated to formulate a cell density for each region. Other organs routinely processed and examined included liver, lung, kidney, heart, skeletal muscle, adipose and haired skin, for which several background findings were noted, but no treatment-related effects were detected.

### Chimaera Generation

Rat blastocysts at E4.5 days post-coitum were collected by noon on the day of injection and cultured for 2–3 hours in KSOM embryo culture medium to ensure cavitation. Cells were disaggregated in TVP, pelleted in N2B27 and pre-plated on gelatin-coated tissue culture plastic in 2iL for 45-60 minutes. Non-attached cells were pelleted and resuspended in N2B27 containing 20 mM HEPES buffer then kept on ice prior to injection. Blastocycts were injected with 10–12 cells, then surgically transferred into the uteri of pseudopregnant Sprague Dawley rats.

### Data availability

The data underlying this report are available in the article (Supplementary Tables S1 and S2).

## Ethics statement

Animal work conformed to guidelines for animal husbandry according to the UK Home Office and approval by the Roslin Institute Animal Ethics Committee. Animals were naturally mated and sacrificed under schedule 1, procedures that do not require specific Home Office approval. Surgical procedures were performed under Procedural Project Licences 6004539 and 6003887 held by M.G.F.S.

## Author Contributions

S.M., K.S-D., and T.B. conceived and designed the study. S.M., K.S-D., L.S and M.G.F.S. performed the study. J.D-P., and D.W. planned and performed the histological analysis. S.M., K.S-D., J.D-P., D.W., and T.B. analysed the data. S.M., and T.B. wrote the manuscript. All authors reviewed the manuscript.

### Acknowledgments

The authors would like to express their gratitude to Mr William Mungall, Mrs Julie Thompson and Mrs Ailsa Travers for their excellent technical support at the Bioresearch and Veterinary Services at the University of Edinburgh; Professor Austin Smith and Dr Kathryn Blair, University of Cambridge for providing the DAK31 cell line and Dr. Tetsushi Sakuma and Professor Takashi Yamamoto, Hiroshima University for their advice and guidance in TALEN construction. This work was supported by funding from the Biotechnology and Biological Sciences Research Council Institute Strategic Programme Grants BB/J004316/1, BB/J004332/1: BBSRC Response mode grant BB/H012478/1, and European Community’s Seventh Framework Programme (FP7/2007-2013) under grant agreement No. HEALTH-F4- 2010-241504 (EURATRANS).

